# Increased α-ketoglutarate as a missing link from the C_3_-C_4_ intermediate state to C_4_ photosynthesis in the genus *Flaveria*

**DOI:** 10.1101/2022.06.01.494409

**Authors:** Qiming Tang, Yuhui Huang, Xiaoxiang Ni, Ming-Ju Amy Lyu, Genyun Chen, Rowan Sage, Xinguang Zhu

## Abstract

As a complex trait, C_4_ photosynthesis has multiple independent origins in evolution. Phylogenetic evidence and theoretical analysis suggest that C_2_ photosynthesis, which is driven by glycine decarboxylation in the bundle sheath cell, may function as a bridge from C_3_ towards C_4_ photosynthesis. However, the exact molecular mechanism underlying the transition between C_2_ photosynthesis towards C_4_ photosynthesis remains elusive. Here, we provide multiple evidence suggesting a role of higher α-ketoglutarate (AKG) concentration during this transition. Metabolomic data of 12 *Flaveria* species, including multiple photosynthetic types, show that AKG concentration initially increases in the C_3_-C_4_ intermediate with a further increase in C_4_ species. Petiole feeding of AKG increased the concentrations of C_4_ related metabolites in C_3_-C_4_ and C_4_ species but not the activity of C_4_ related enzymes. Sequence analysis shows that glutamate synthase (Fd-GOGAT), which catalyzes the generation of glutamate using AKG, was under strong positive selection during the evolution of C_4_ photosynthesis. Simulations with a constraint-based model for C_3_-C_4_ intermediate further show that decreasing the activity of Fd-GOGAT facilitates the transition from a C_2_-dominant to a C_4_-dominant CO_2_ concentrating mechanisms. All these provide an insight into the mechanistic switch from C_3_-C_4_ intermediate to C_4_ photosynthesis.

## INTRODUCTION

Plants with C_4_ photosynthesis have a higher light energy conversion efficiency, higher water, and nitrogen use efficiencies compared with C_3_ plants(Dengler and Nelson, 1999; Vogan and Sage, 2011; Zhu, et al., 2010). C_4_ plants also generally grow faster than C_3_ plants, which contribute 23% of primary productivity on land with 3% of the species in the plant kingdom (Sage, et al., 2012). In many of over 60 lineages, species are known to be both phylogenetically intermediate and both physiologically and anatomically intermediate between the full C_3_ and C_4_ character states (Araus, et al., 1990; Kennedy and Laetsch, 1974; McKown and Dengler, 2007; Monson, et al., 1989; Monson, et al., 1984). These intermediates have formed the basis for studies that evaluate how C_4_ photosynthesis has repeatedly evolved, as well as addressing the larger question of how complex traits can evolve with high frequency. While a general model of C_4_ evolution has been developed (Heckmann, et al., 2013; Sage, et al., 2012), key steps in the process remain uncertain, notably, how the C_4_ metabolic pathway arose from a background of modifications termed C_2_ photosynthesis. In C_2_ photosynthesis, CO_2_ released from glycine decarboxylase (GDC) in bundle sheath cells (BSC) provides the substrate for RuBisCO carboxylation. On one hand, the C_2_ photosynthesis can confer lower CO_2_ concentrating point and increased competitive advantage over C_3_ species under stress conditions (Monson, 1989; Vogan and Sage, 2012); on the other hand, the GDC in BSC can create an imbalance in the ammonia residue between BSC and mesophyll cell (MC) (Bräutigam and Gowik, 2016; Rawsthorne, et al., 1988). Various hypothetical pathways were proposed to address the issue of ammonia misbalance between BSC and MC (de Oliveira Dal’Molin, et al., 2010; Mallmann, et al., 2014; Monson and Rawsthorne, 2000). Among these solutions, a C_4_/C_2_ mixed pathway which includes major steps of the C_4_ photosynthetic pathways (Mallmann, et al., 2014), has been proposed and regarded as a bridge between C_2_ and C_4_. However, so far, how C_4_ metabolism emerges from intermediate photosynthesis remains elusive.

This study aims to address how C_4_ photosynthesis emerged from C_3_-C_4_ intermediate photosynthesis. To study this question, we used the genus *Flaveria* as our model plant system. We made this choice since not only the genus of *Flaveria* is one of the youngest known C_4_ lineages (Sage, et al., 2012), but also various aspects related to C_4_ photosynthesis evolution have been well characterized in the genus *Flaveria*, including various physiological indicators (Brown, et al., 1986; McKown, et al., 2005), changes in the transcriptomics (Lyu, et al., 2021; Lyu, et al., 2015; McKown and Dengler, 2007), and activities of key C_4_ enzymes (Ludwig, 2011; McKown and Dengler, 2007) and metabolite concentrations related to the C_2_ and C_4_ photosynthesis (Borghi, et al., 2021; Gowik, et al., 2011; Ubierna, et al., 2013).

Specifically, in this study, we identified the content of α-ketoglutarate (AKG) whose changes are most relevant to C_4_ evolution. The comparison combined with the impacts of external application of AKG on different *Flaveria* species and model simulations show a major role of increased content of AKG during the shift from C_2_ to C_4_ metabolism, hence fills in a major missing link during the evolution from C_3_-C_4_ intermediate to C_4_ photosynthesis.

## RESULTS

### Generation of *Flaveria* metabolic and transcriptomic profiles

To map the changes of metabolism along C_4_ evolution, we conducted primary metabolic profiling and transcriptomic sequencing for 12 *Flaveria* species, including 2 species categorized as C_3_ photosynthesis: *F. cronquistii* (*F. cro*) and *F. robusta* (*F. rob*), 2 C_4_ photosynthesis: *F. australasica* (*F. aus*) and *F. bidentis* (*F. bid*), 3 C_4_-like: *F. brownii* (*F. bro*), *F. vaginata* (*F. vag*) and *F. palmeri* (*F. pal*) and 5 C_3_-C_4_ intermediate species: *F. sonorensis* (*F. son*), *F. angustifolia* (*F. ang*), *F. ramosissima* (*F. ram*), *F. chloraefolia* (*F. chl*) *and F. linearis* (*F. lin*), with 4 species from the base of the phylogeny tree, 5 species from the A clade and 3 species from the B clade of the phylogeny tree (Fig.1A) (Lyu, et al., 2015).

**Figure 1.**
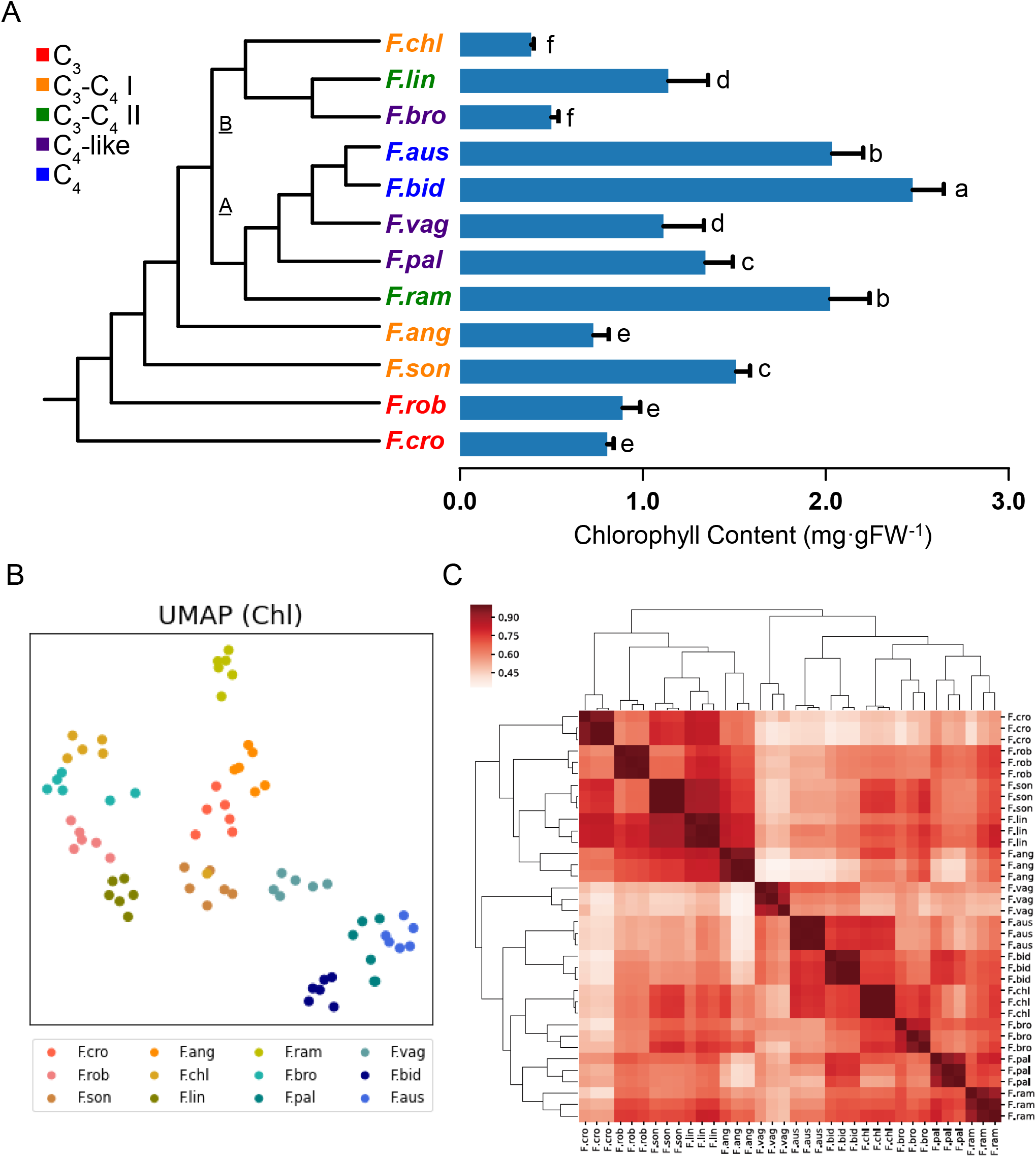
Overview of *Flaveria* metabolic and transcriptomic profiles. A) Phylogenetic tree and chlorophyll content of 12 flaveria species sampled. Different letters indicate a significant difference between the medians (Kruskal-Wallis tests, α = 0.05, p-values adjusted using false discovery rate) (Benjamini and Hochberg, 1995). n = 5. B) Dimensionality reduction analysis (UMAP) of metabolomics data normalized on chlorophyll content. Data were normalized using Z-score. C) Heatmap of Pearson correlation coefficient among transcriptomics samples. Duplicate species names represent different biological samples.

On the metabolic level, we quantified the concentrations of forty metabolites involved in the primary metabolism using HPLC-MS/MS. The profile covers metabolites involved in the Calvin-Benson cycle, glycolysis, TCA cycle, photorespiration, nitrogen assimilation and several metabolites involved in energy and redox metabolism. The chlorophyll content of each species was significantly different (Fig 1A), the metabolic variations among species could be different when the calculation is based on fresh weight or chlorophyll content. In our case, the concentrations of all metabolites were normalized on total chlorophyll content basis. Almost all biological repeats are clustered together in UMAP plot, which indicated the reliability of metabolomic samples (Fig 1B).

On the transcriptomic level, we obtained on average 25.96 million raw reads for each sample. From 65,144 to 163,967 transcripts were *de novo* assembled based on raw reads for each species. On average, 60.1% of those transcripts were functionally annotated based on the protein reference from *Arabidopsis thaliana*. To quantify the transcript abundance of the *Flaveria* species, raw reads were mapped to assembled transcripts of corresponding species, which resulted in mapping rates percentages ranging from 73.49% to 96.43% for each sample (Table. S1). Transcriptomics profiles of all *Flaveria* species showed high Pearson correlation coefficients ranging from 0.85 to 1.00 between biological replicates, suggesting the reliability of transcriptomics data. Besides, the tree topology of the twelve species based on correlations of transcriptomic abundances is largely consistent with that of the phylogenetic tree based on coding sequences on transcriptomic level (Fig 1C). This consistency suggested a co-evolution of gene sequence and transcript abundance as reported recently (Lyu, et al., 2021).

### AKG stepwise increased in the evolution of C_4_ photosynthesis in *Flaveria*

We calculated the importance weight of each metabolite in distinguishment of different photosynthetic types by feature selection analysis (see Methods). Specifically, we characterized the 12 *Flaveria* species into different groups. We tested 4 different schemes of priori grouping (Fig S1A). Among the metabolites resulted in a cumulative importance of 80% (Fig S1B) in each priori grouping, five of them are shared, i.e., α-ketoglutarate (AKG), Ala, Ser, citrate (CIT) and fructose diphosphate (FBP) (Fig S2). In these 4 priori grouping schemes, AKG has always been the top in importance rank (Fig 2B S1B). This result suggests that the content of AKG reflected the evolutionary trajectory of C_4_ photosynthesis in the genus of *Flaveria*.

**Figure 2.**
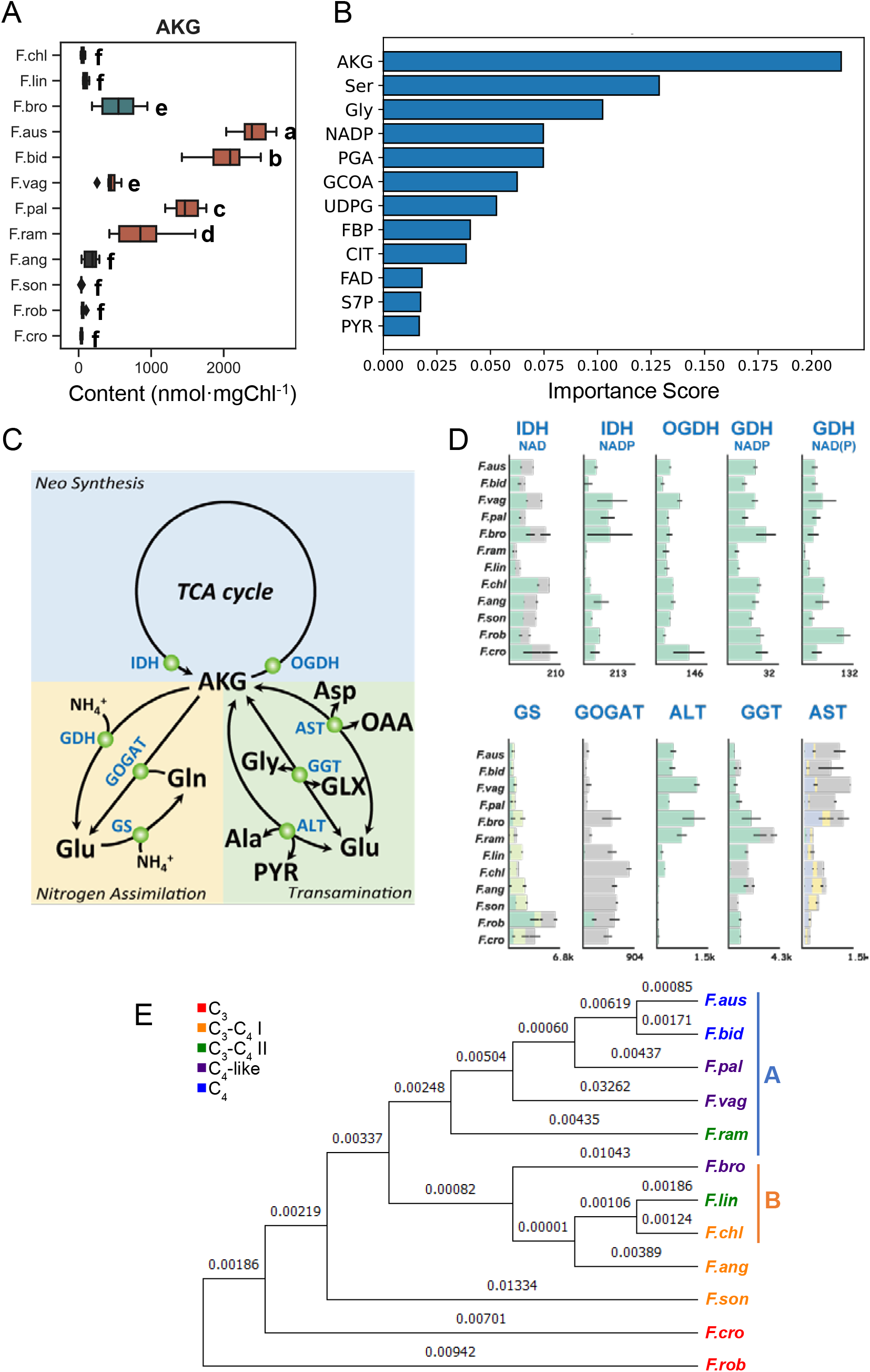
A) Box plot of AKG content. Box with black background indicate the ancestor species in phylogenetic tree, orange indicate the species belong clade A in phylogenetic tree, and green indicate the species belong clade B in phylogenetic tree. Different letters indicate a significant difference between the medians (Kruskal-Wallis tests, α = 0.05, p-values adjusted using false discovery rate) (Benjamini and Hochberg, 1995). n = 6. B) important metabolites in scheme #1 with a cutoff of 80% cumulative importance (See Fig S1 for other schemes). C) Metabolic pathways related to AKG formation and metabolism. D) AKG-related gene expression (TPM) plots for all expressed homologous genes annotated with same E.C. number. Error bar indicates S.D., n = 3. The maximum total expression value in all species is marked in the lower right. E) Phylogenetic tree of Fd-GOGAT (AT5G04140) protein sequence. The value above the branch represents the length of the phylogenetic branch. A and B represent the species belonging to clade A and clade B on the phylogenetic tree of Flaveria respectively.

Examination of AKG in different species shows a stepwise increase of AKG concentration along the C_4_ evolution in the genus of *Flaveria* (Fig 2A, S2). The first step occurred at *F. ram*, a type II C_3_-C_4_ intermediate with an undeveloped C_4_ CCM. Second step occurred at *F. bid*, a model C_4_ species. That is, AKG increased at two key nodes of C_4_ evolution, i.e., the first key node co-occurred with the preliminary C_4_ metabolism in *F. ram*, while the second key node co-occurred with the complete C4 cycle (*F. bid*). These results suggest that the gradual increase of AKG content may have a physiological significance during the evolution of C_4_ photosynthesis.

We further examined what might be a direct cause of changes in AKG content. All AKG-related primary metabolic pathways are summarized in Fig. 2C, which include neo-synthesis of AKG (IDH and OGDH), nitrogen assimilation (GS-GOGAT and GDH) and transamination related reactions (AST, ALT and GGT). The expression of four enzymes (AST, ALT, GS and Fd-GOGAT) shows a regular pattern in the evolution of C4 photosynthesis (Fig 2D). Though the expression of AST and ALT were increased, however, their increased expression levels do not necessarily increase AKG concentrations since these two enzymes catalyze reversible reaction. Besides, the expression levels of GS and Fd-GOGAT were evidently lower in C_4_ species (Fig. 2D). Further, we found that the protein sequence of Fd-GOGAT was under positive selection along the evolution of C_4_ photosynthesis (*p* = 7.49×10^−5^, see Methods), and the phylogenetic tree of Fd-GOGAT protein sequence is very similar to that of species (Fig 2E). All these suggest that the decreased Fd-GOGAT might have been a primary reason for the increased concentration of AKG in the C_4_ species.

### AKG promotes photosynthetic efficiency in developing C_4_ photosynthesis

Given the increase of AKG concentration along C_4_ evolution, we speculate that there is a positive physiological role of AKG on photosynthesis of leaves from intermediate species, at which stage when the AKG concentration initially increased. We tested this using exogenous feeding of AKG through petiole. As the species where AKG initially increased during the evolution, *F. ram* is the first to test with AKG, sorbitol (as osmic control) and Ala (another shared important metabolite in feature selection analysis, and same increased during evolution with AKG). Results show that exogenous AKG feeding indeed showed major impacts on photosynthetic properties of intermediate compared with Ala (Fig. 3). Specifically, AKG-treated *F. ram* showed a higher net photosynthesis rate compared with osmotic control, but Ala-treated *F. ram* not. Furthermore, AKG-treated *F. vag* and AKG-treated *F. aus* both showed a positive effect in photosynthesis rate (Fig. 3). All three of them belong to clade A in the phylogenetic tree. On the contrary, as a base species from phylogeny the photosynthesis rate of AKG-treated *F. ang* did not changed. In *F. bro*, a B branch C_4_-like *Flaveria* species, the AKG treatment did not increase, rather decreased, the photosynthetic rate (Fig. 3), suggesting that the metabolism of the B branch of *Flaveria* might have different evolutionary trajectory compared to the A branch.

**Figure 3.**
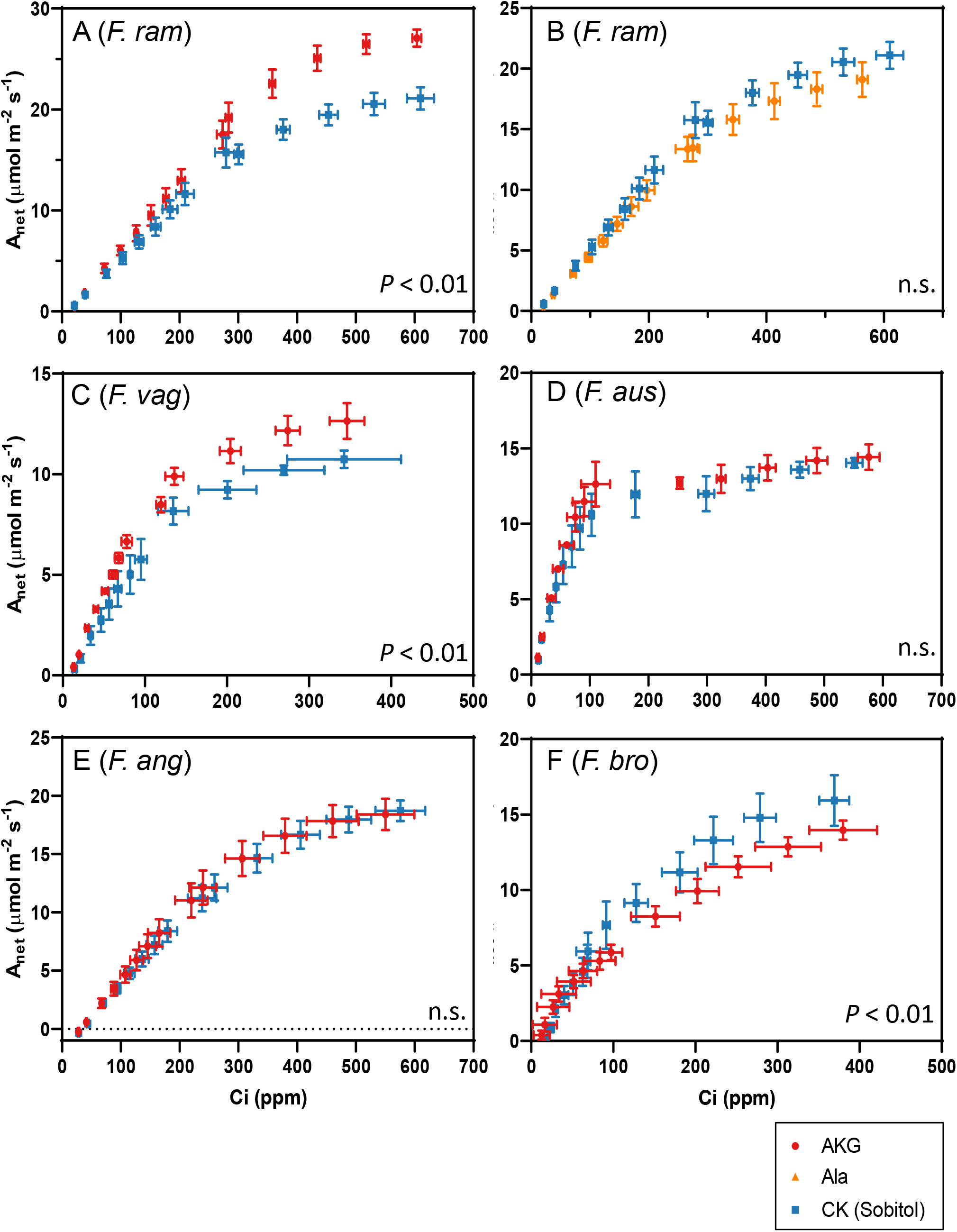
Exogenous AKG showed differential effects on Flaveria species with different photosynthetic types. A-F) CO_2_ response curves of Flaveria species treated with metabolites. the data points were the average values of 3-4 biological samples in each group, and the error bar was the standard error (S.E.). Significant differences between treatments were identified using contrasts analysis (lsmeans package, R), n.s.: not significant.

Given the positive impact of AKG on leaf photosynthetic rate, we further tested the effects of AKG treatment on the metabolomics of these five species of *Flaveria*. We found that, except AKG itself, SUC and FUM generally increased, which indicate a higher respiration activity in all AKG-treated *Flaveria* (Fig. 4A). But the content of PEP, OAA, MAL and PYR increased in AKG-treated *F. ram* and OAA, MAL and PYR increased in AKG-treated *F. vag* (Fig. 4A). Considering the higher net photosynthesis rate under AKG-treated *F. ram* and AKG-treated *F. vag* (Fig 3), we suggest that, in AKG-treated C_3_-C_4_ intermediate of *Flaveria*, the efficiency of C_4_ photosynthesis was improved and covered the higher respiration activity.

**Figure 4.**
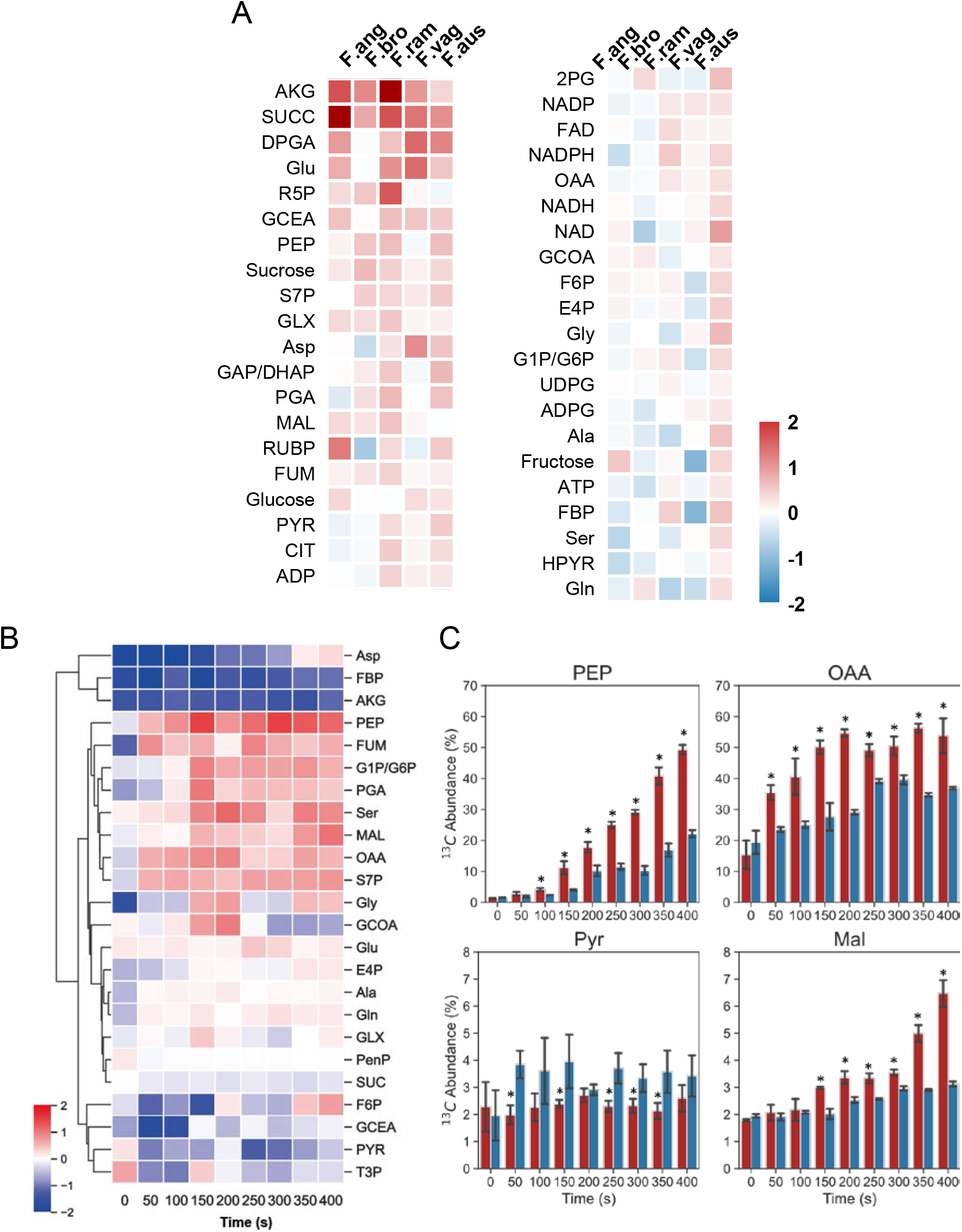
Exogenous AKG promotes the C4 photosynthetic flux. A) Changes in leaf metabolite concentrations for Flaveria species treated with different metabolites (values in heatmap indicate the log2 fold changes compared to the control group). B,C) 13C abundance change of F. ram treated with AKG. B) values in heatmap indicated the log2 fold change compared to control group. C) Histogram plot for four C4-related metabolites. error bar indicates S.D., *: p < 0.05 (Student’s t-test). n = 4.

^13^CO_2_ labelling experiment was used to further test whether C_4_ flux was increased in AKG-treated *F. ram*. The result shows that the accumulation rate of ^13^C abundance in PEP, OAA and MAL was significantly faster in AKG treated *F. ram* compared to osmic control group, which indicates a faster turnover rate of PEPC and NADP-MDH (Fig. 4B, C). While the enzyme activities of PEPC and NADP-MDH did not increase under AKG treatment (Fig. S3), which concludes that the AKG promotes C_4_ photosynthetic flux by metabolic content rather than the enzyme activity.

### A constrained flux of Fd-GOGAT promotes the shifting from C_2_ to C_4_ CCM in model simulation

Given that the decreased expression of Fd-GOGAT along C_4_ evolution and also the strong positive selection signal in this enzyme in C_4_ species, we tested whether Fd-GOGAT can directly affect C_4_ photosynthesis using a mechanistic C_3_-C_4_ intermediate model, which includes both C_2_ CCM and C_4_ CCM. Specifically, we parameterized the C_3_-C_4_ photosynthesis model (Mallmann, et al., 2014) with physiological and transcriptomics data of type II C_3_-C_4_ *Flaveria* species and then conducted flux balance analysis. In our simulation study, serval constrains in the original model were modified: a) we kept the upper bound of RubisCO Oxygenation rate and PEPC carboxylation rate but changed their lower bound to 0; b) we constrained the net AKG generation rate to a constant value and c) we constrained the synthesis of glutamate only from Fd-GOGAT (Fig 5A, See methods for detail). The simulation results show that when the flux of Fd-GOGAT decreases, the flux of GDC in BSC decreased, while the flux of NADP-ME in BSC increased when the flux of Fd-GOGAT gradually decreases (Fig. 5B). Our simulation shows that there could be a “switch point” where the system is switched from C_2_-dominant to C_4_-dominant metabolic pathways (Fig. 5B). When the system near this switch point, multiple amino acids and malate are transported in plasmodesmata to optimize ammonia balancing and C concentrating (Fig 5C). Since AKG is the substrate of Fd-GOGAT, this simulation results are consistent with a higher AKG concentration and a lower GS-GOGAT expression level in *F. ram* and C_4_ species of *Flaveria*. These results suggests that a decreased activity of Fd-GOGAT may increase the concentration of AKG, facilitating the switch from a C_2_-dominant metabolism to C_4_-dominant metabolism.

**Figure 5.**
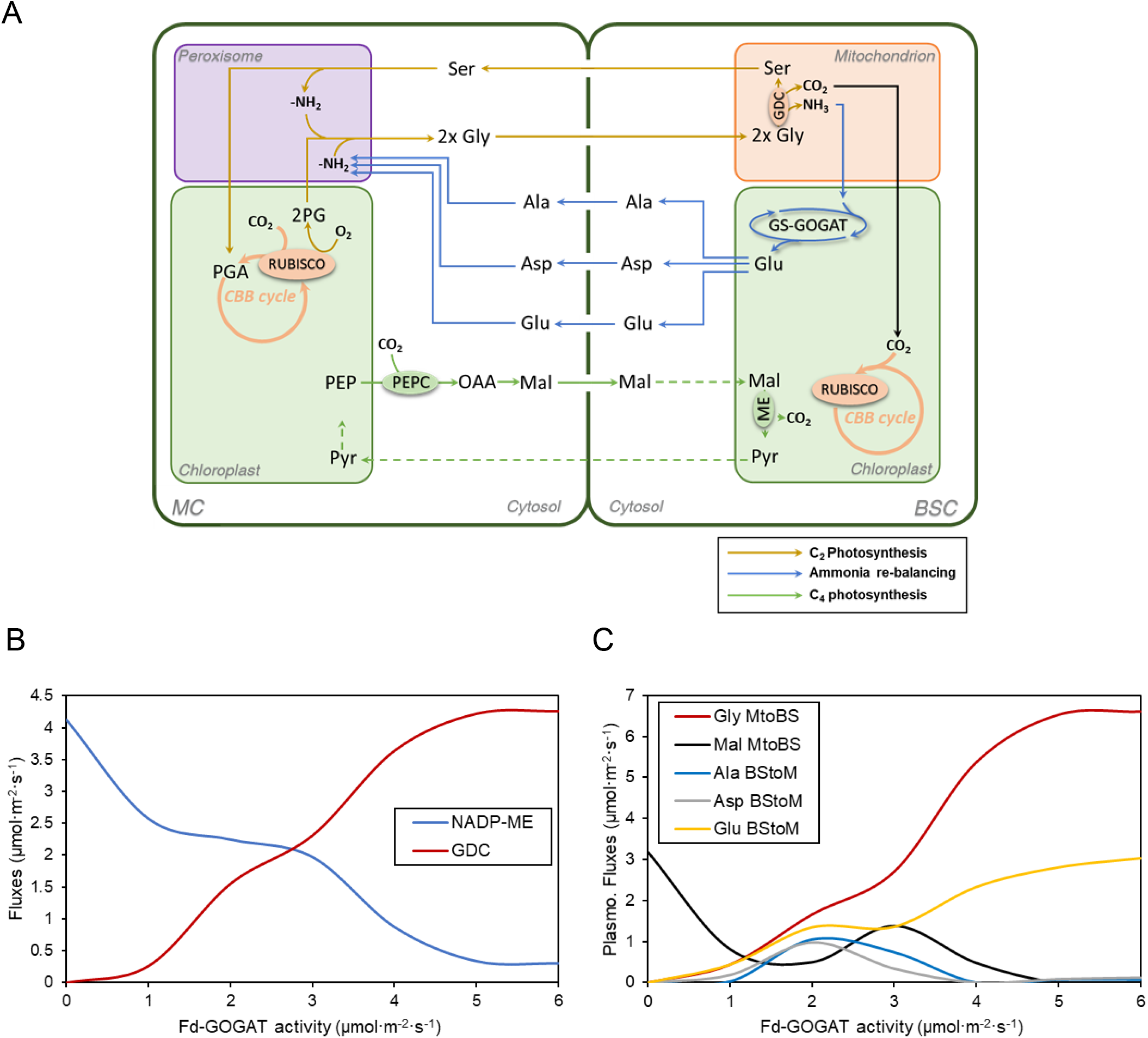
Model simulations suggest that a low Fd-GOGAT activity facilitates the shifting from a C_2_ to C_4_ CCM in model simulations. A) Mechanistic switching from C_2_/C_4_ mixed photosynthesis to C_4_ photosynthesis when GS-GOGAT activity is limited. B) Predicted fluxes of NADP-ME and GDC in bundle sheath cells at different activities of Fd-GOGAT. C) Predicted plasmodesmata transfer fluxes of amnio acids and malate between mesophyll cells and bundle sheath cells at different activities of Fd-GOGAT. MtoBS: flux transfer from mesophyll cells to bundle sheath cells; BStoM: flux transfer from bundle sheath cells to mesophyll cells.

## Discussions

This study provides multiple evidence supporting an important role of increased AKG concentration during the formation of a C_4_ metabolism from C_2_ photosynthesis. First, we found that there is a dramatic increase in the concentration of AKG along C_4_ evolution on chlorophyll content basis. In previous study, the content changes of AKG in different photosynthetic types of *Flaveria* were also be reported with relative abundance (Gowik, et al., 2011), or fresh weight basis (Borghi, et al., 2021). In (Gowik, et al., 2011), AKG increased in C_3_-C_4_ intermediate relative to C_3_, but (Borghi, et al., 2021) not, which could be a more humid condition and large temperature difference between day and night in (Borghi, et al., 2021). Second, under AKG-treatment, we found that the concentrations of critical metabolites involved in C_4_ photosynthesis, i.e. OAA and PYR, increased (Fig 4A) and the leaf net photosynthetic rate increased (Fig 3A). Finally, Fd-GOGAT showed the greatest number of amino acid residues which are under positive selection (*p* = 7.49×10^−5^). Considering that the decreased Fd-GOGAT activity can in principle result in an increased concentration of AKG, the increased AKG concentration should be closely linked to the C_4_ evolution.

Increase in the AKG concentration can directly increase the C_4_ flux. AKG can affect the direction and also reaction speed of transamination reactions involving the conversion between AKG and glutamate. At 25°C (1 Standard atmospheric pressure (atm)) aqueous solution, the change in Gibbs free energy of aspartate transaminase (AKG + Asp ⇌ Glu + OAA, EC: 2.6.1.1) is 2.8 ± 1.1 kJ/mol (pH: 7.0) when the ratio of reactant concentrations is 1:1:1:1. However, if the abundance of AKG increased 5 times, i.e., the ratio of reactant concentrations is 5:1:1:1, the change in Gibbs free energy would be -1.1 ± 1.1 kJ/mol (pH: 7.0). Reaction would proceed in the direction of generating OAA. Therefore, the relative concentration of AKG can influence the direction of the flux of aspartate transaminase, and same with alanine aminotransferase (EC: 2.6.1.2). High concentration of AKG helps maintain the high concentration of C_4_ related metabolites (PYR and OAA), which promotes C_4_ flux, as is clearly shown by the increased net photosynthesis rates when *F*.*ram* was fed with AKG through petiole (Fig 3A).

Besides a direct role of AKG in the C_4_ metabolism, AKG may also influence fluxes through the C_2_ pathway. AKG is also involve in the transamination reactions in the C_2_ pathway (GGT, EC: 2.6.1.4) and GS-GOGAT system, decreasing the AKG concentration will decrease the flux through these reactions. Furthermore, the flux of C_2_ CCM flux are typically influenced by a number of factors, which include the RuBP oxygenation catalyzed by RubisCO, the formation of glycine shuttle between MC and BSC, and finally the activities of enzymes, such as Fd-GOGAT. Given that the AKG can influence reactions in both the C_2_ pathway and the C_4_ pathway, we explicitly tested this idea with a mechanistic model with C_3_-C_4_ intermediate photosynthesis. Our simulation indicates that there was a switch from C_2_ to C_4_ photosynthesis when the concentration of AKG is increased to a certain level (Fig. 5B). In fact, after an efficient C_4_ photosynthesis is established, carboxylation of RubisCO in mesophyll cells can no longer be the dominant carbon assimilation mechanism, i.e. it could be gradually diminished during the evolution. In line with this, in our earlier analysis of C_3_-C_4_ intermediate, i.e. a metabolic state where the C_2_ and C_4_ co-exist, we have shown that there is little or no competitive advantage in terms of photosynthesis per leaf area (Wang, et al., 2017).

## Materials and methods

### Plant material

Seeds or plants materials for *Flaveria* species used were kindly provided by Prof. Rowan Sage (University of Toronto, Canada). The plants used for sampling were grown from cuttings and grown in the greenhouse. Growth condition of the greenhouse followed our previous work (Lyu, et al., 2015), i.e. growth temperature: 28 °C; photoperiod: 14h light / 10h dark; relative humidity (RH): 60-65% and photosynthetic photon flux density (PPFD): 500 μmol m^−2^ s^−1^.

### Metabolite treatments and gas exchange measurements

To feed specific metabolites to plants, root was cut and the bottom part of the stem was soaked in 10mM solution of the corresponding metabolite separately. 10 mM sorbitol solution was used as the control group. All solutions’ pH has adjusted to 6.5. Plants were illuminated under light with a photosynthetic photon flux density (PPFD) of 400 μmol·m^−2^·s^−1^ for 1 hour before gas exchange measurement. Leaves were sampled 1 hour after light treatments.

The LI-6800 Portable Photosynthesis System (Li-Cor, Inc., Lincoln, NE, USA) was used for gas exchange measurements. For the CO_2_ response curve measurements, air flow rate was set to be 500 μmol/s, block temperature of the chamber was set to be 28 °C, reference CO_2_ concentration was set to be 400 ppm and the PPFD was set to be 1800 μmol·m^−2^·s^−1^. Plants were maintained under such conditions for 15 mins before the first measurement. In the first loop, the reference CO_2_ concentration was set to the following sequence of CO_2_ concentrations, i.e. 400, 300, 260, 220, 180, 140, 100, 50 and 25 ppm; at each CO_2_ level, leaves were maintained for 2 mins before data logging. After this sequence of CO_2_ concentrations, the reference CO_2_ concentration was set back to 400 ppm and maintained for 15 mins before the second loop began. In the second loop, the reference CO_2_ concentrations was set in the following order: 400, 500, 600, 700, 800ppm with plants maintained at each concentration for 2 mins before logging. The reference CO_2_ concentration was set to 400 ppm when the second loop is finished.

### LC-MS/MS and Metabolomics analysis

The LC-MS/MS analysis was done following (Arrivault, et al., 2019; Wang, et al., 2014). Similar to the sampling procedure for transcriptomics analysis, the newly fully expanded and illuminated leaves, usually the 2^nd^ or the 3^rd^ pair of leaves counted from the top of each branch, were used for all species. For each sample, a leaf sample with a 2 cm^2^ leaf area was sampled and frozen in liquid nitrogen instantaneously. For some species which have needle-like leaves (*F. ram* and *F. chl*), two leaves were pieced together. Several leaves with same leaf position were sampled to measure the fresh weight and chlorophyll concentration, which were used to convert the concentration of metabolites between fresh weight and chlorophyll concentration basis.

All leaf samples were cut *in situ* and immediately transferred into a pre-frozen 2mL EP tube, then stored in liquid nitrogen for extraction. After grinding, each sample was fully dissolved with 800 μL extraction buffer (methanol: chloroform = 7:3 (v/v), -20°C pre-cooling) and incubated under -20°C for 3 hours. Then 560 μL distilled water (ddH_2_O) was add and mixed with each sample, 800 μL supernatant was extracted after centrifugation (×2200g, 10min, 4°C). After that, 800ul buffer (methanol: ddH2O = 1: 1(v/v), 4°C pre-cooling) was mixed with the sample for another extraction. For each sample, 1.6 mL supernatant in total was obtained by filtering the extraction buffer with a 0.2 μM nylon filter. Among them, 1 mL was used for MS/MS analysis and 20 uL was used for QC sample. All extraction operations were performed on ice.

Luna NH_2_ column (3μm, 100mm*2mm, Phenomenex co. Ltd, USA) was used in the liquid chromatography. The LC gradient was set with eluent A, which has 10 mM Ammonium acetate and 5%(v/v) acetonitrile solution, with the pH adjusted to 9.5 using ammonia water and eluent B (acetonitrile): 0-1 min, 15% A; 1-8 min, 15-70% A; 8-20 min, 70-95% A; 20-22 min, 95% A; 22-25 min, 15% A. In the aspect of mass spectrometry analysis, QTRAP 6500+ (AB Sciex, co. Ltd, USA) was used in MRM model, all parameters used following (Arrivault, et al., 2019; Wang, et al., 2014) with optimization (Supply Dataset 1). Concentration of all metabolites in samples were calculated based on the “concentration-peak area” curve of standard samples and converted to nmol·gFW-1 with specific leaf weight of each species measured before.

### ^13^CO_2_ labelling

^13^CO_2_ labelling of AKG-treated *F. ram* followed (Heise, et al., 2014). All treated individuals were pre-illuminated at a PPFD of ∼500 μmol·m^−2^·s^−1^ for 30 mins before labelling. During the labelling, the CO_2_ concentration was kept at 450 ppm, relative humidity (RH) was kept at 40-60%. When ^13^C labelling finished, leaf sample was transferred into liquid nitrogen immediately. Then about 30 mg of frozen leaf tissue was used for metabolite extraction. The extraction protocol used is same as earlier used for metabolite profiling with parameters for mass spectrometry optimized (Supply Dataset 1).

### Transcriptome assembly and quantification

To obtain the transcriptomics data, we sampled the newly developed fully expanded leaves, usually the 2^nd^ or the 3^rd^ pair of leaves counted from the top for each species. The chosen leaves were cut and immediately put into liquid nitrogen and kept in -80 °C liquid nitrogen before further processing. Total RNA was isolated following the protocol of the PureLInkTM RNA kit (Thermo Fisher Scientific, USA). Illumina sequencing was performed and analysis were conducted following our previous work (Lyu, et al., 2021). Specifically, transcript abundances of *Flaveria* samples were calculated as Transcripts Per kilobase of exon model per Million mapped reads (TPM) using the RSEM package (version 1.3.1)(Li and Dewey, 2011).

### Enzyme activity measurements

The PEPC activity was assayed following (Fukayama, et al., 2003). The NADP-ME activity and the NADP-MDH activities were assayed followed (Tsuchida, et al., 2001). The PPDK activity was assayed following (Wang, et al., 2008). CARY50 UV Spectrophotometer (VARIAN Co. Ltd, USA) was used to monitor the consumption or generation of NAD(P)H at 340 nm.

### Feature selection analysis

Feature selection analysis was performed with a python script (https://github.com/WillKoehrsen/feature-selector). The method finds features with a gradient boosting machine implemented in the LightGBM library (https://github.com/Microsoft/LightGBM). Analysis implemented with default parameters and the metric used for early stopping is set to the error rate for mulit-class classification (Ke, et al., 2017; Li, et al., 2007).

### Identification of amino acid residues under positive selection

To identify positively selected amino acid residues in Fd-GOGAT in *Flaveria* C_4_ species, we used *Flaveria* non-C_4_ species (including C_3_ and C_3_-C_4_ species) as background. For each gene, only these genes with completely assembled sequence were included in the analysis. To do this, firstly, protein sequences of orthologous genes were aligned using the software Muscle (Edgar, 2004). Then aligned protein sequences were used to guide the codon-wise alignment of CDS with PAL2NAL (Suyama, et al., 2006). After the gaps and stop codons were removed, multiple sequence alignment results were input into the PAML software (version 4.8) for positive selection tests (Yang, 2007). In this study, the positive selection test was conducted with the branch-site model (model=2, NSsites=2) under the nucleotide substitution conditions of CodonFreq=2 (4 distinct frequencies are used for each position, named F3×4). The likelihood of the null hypothesis was calculated under this branch-site model with fixed dN/dS ratio (ω=1, neutral). The maximum likelihood of the alternative hypothesis was calculated under this branch-site model with flexible dN/dS ratio (ω>1, positive selection). Then, the likelihood ratio test (LRT) was conducted between the null hypothesis and the alternative hypothesis under chi-square distribution. Those genes identified to be significant (p-value < 0.05, Benjamini Hochberg (BH) adjusted) by tests under these two different nucleotide substitution parameter settings were considered as genes under positive selection.

### Simulation of C_2_ and C_4_ Photosynthesis with a constrained Fd-GOGAT flux

A previously reported genome-scale constraint based model of C3–C4 photosynthesis (Mallmann, et al., 2014) was used to test the impacts of different Fd-GOGAT activities on metabolic fluxes. We parameterized this model with parameters for a type II C3-C4 state as given in (Heckmann, et al., 2013), which the flux of C4 CCM and flux of C2 CCM are incorporated together. In the original model, the flux of RubisCO Oxygenase and PEPC are constrained to experimental values from (Heckmann, et al., 2013). In our analysis, these fluxes were set to be flexible while the flux of Fd-GOGAT was set to be under limitation. So, we kept the upper bounds of RubisCO oxygenation rate and PEPC carboxylation rate but changed the lower bound of these two fluxes to be zero. Furthermore, we set the respiration rate to be constant (Byrd, et al., 1992), i.e. the net AKG generation rate (IDH) is set as constant to ensure the AKG generation rate would not be a limitation of model (Mallmann, et al., 2014). Finally, we constrain the synthesis of glutamate can only be carried out through Fd-GOGAT, which is driven by photosynthetic reducing energy.

## ABBREVIATION

**Species:**

F. ang: Flaveria angustifolia
F. aus: Flaveria australasica
F. bid: Flaveria bidentis
F. bro: Flaveria brownii
F. chl: Flaveria chloraefolia
F. cro: Flaveria cronquistii
F. lin: Flaveria linearis
F. pal: Flaveria palmeri
F. ram: Flaveria ramosissima
F. rob: Flaveria robusta
F. son: Flaveria sonorensis
F. vag: Flaveria vaginata

**Metabolite:**

2PG: 2-phosphoglycolate
AKG: α-ketoglutarate (2-oxoglutarate)
Ala: Alanine
CIT: Citrate
E4P: Erythrose-4-phosphate
F6P: Fructose-6-phosphate
FBP: Fructose bisphosphatase
Fd: Ferredoxin
FUM: Fumarate
GCEA: Glycerate
GCOA: Glycolate
Gln: Glutamine
Glu: Glutamate
GLX: Glyoxylate
Gly: Glycine
HPYR: Hydroxypyruvate
MAL: Malate
OAA: Oxaloacetate
PenP: Pentose phosphate
PEP: Phosphoenol pyruvate
PGA: Phosphoglycerate
PYR: Pyruvate
RuBP: Ribulose 1,5-bisphosphate
S7P: Sedoeheptulose-7-phosphate
Ser: Serine
SUC: Succinate
T3P: Triose phosphate

**Gene/Enzyme:**

ALT: Alanine transaminase [EC:2.6.1.2]
AST: Aspartate aminotransferase [EC:2.6.1.1]
Fd-GOGAT: Ferredoxin-dependent glutamate synthase [EC: 1.4.7.1]
GDH: Glutamate dehydrogenase [EC:1.4.1.3 1.4.1.4]
GGT: Glutamate: glyoxylate aminotransferase [EC:2.6.1.4]
GS: Glutamine synthetase [EC: 6.3.1.2]
IDH: Isocitrate dehydrogenase [EC: 1.1.1.41 1.1.1.42]
OGDH: 2-Oxoglutarate dehydrogenase [EC:1.2.4.2]
RubisCO: Ribulose-1,5-bisphosphate carboxylase/oxygenase [EC:4.1.1.39]
SUDH: Succinate dehydrogenase

## Acknowledgement

We thank Yongyao Zhao for the support on cultivation of *Flaveria*. We thank Faming Chen for the valuable suggestions on bioinformatics analysis. We thank Wenli Hu, Xiaoyan Xu and Shanshan Wang of the public technology platform in Institute of Plant Physiology and Ecology (Shanghai, China) for the assistance on metabolomics measurement.

## Accession Numbers

The RNA-seq is submitted to Sequence Read Archive (SRA) in the Nation Center for Biotechnology Information (NCBI) database under the following accession number: PRJNA820135

## Author Contribution

QT and XZ designed the study, performed the analysis, and wrote the paper; QT, YH, XN, ML, GC conducted the experiments and performed the analysis. RFS provided many of the species and assisted with manuscript preparation.

## Supplemental Data

Supplemental DataSet1. MS parameters for metabolic profiling.

Supplemental DataSet2. Metabolic profiles of 12 *Flaveria* species

Supplemental DataSet3. Transcript profiles of 12 *Flaveria* species

Supplemental DataSet4. Source R code used for the simulation with constrain-based models

---

**Table S1.** Statistics of transcriptomics data

| Species | Clean Base (GB) | Assembled Transcripts | Mapping Ratio (%) |
| --- | --- | --- | --- |
| F.ang | 6.7 / 7.94 / 6.94 | 75,861 | 82.00 / 82.72 / 83.01 |
| F.aus | 9.49 / 7.53 / 6.78 | 73,853 | 83.60 / 83.40 / 83.12 |
| F.bid | 7.62 / 8.34 / 7.85 | 65,144 | 86.58 / 86.43 / 85.05 |
| F.bro | 7.36 / 9.34 / 10.29 | 85,926 | 84.12 / 84.06 / 83.13 |
| F.chl | 6.04 / 5.16 / 7.58 | 77,447 | 83.07 / 82.68 / 83.02 |
| F.cro | 6.74 / 8.3 / 6.79 | 91,918 | 77.66 / 77.59 / 79.23 |
| F.lin | 7.1 / 9.13 / 6.63 | 110,727 | 84.59 / 84.68 / 84.94 |
| F.pal | 7.11 / 10.81 / 7.35 | 88,628 | 84.82 / 73.49 / 80.70 |
| F.ram | 6.37 / 6.25 / 7.6 | 163,967 | 81.03 / 81.58 / 81.65 |
| F.rob | 6.95 / 6.23 / 7.1 | 80,514 | 80.01 / 82.41 / 82.95 |
| F.son | 7.06 / 7.03 / 6.73 | 74,137 | 80.73 / 81.26 / 82.14 |
| F.vag | 6.88 / 7.49 / 10.5 | 80,855 | 84.31 / 85.21 / 84.17 |

For each species, clean bases and mapping ratio are counted for three biological samples and distinguished with a slash in table. Transcripts were assembled with pooled reads from all three biological samples

**Figure S1.**
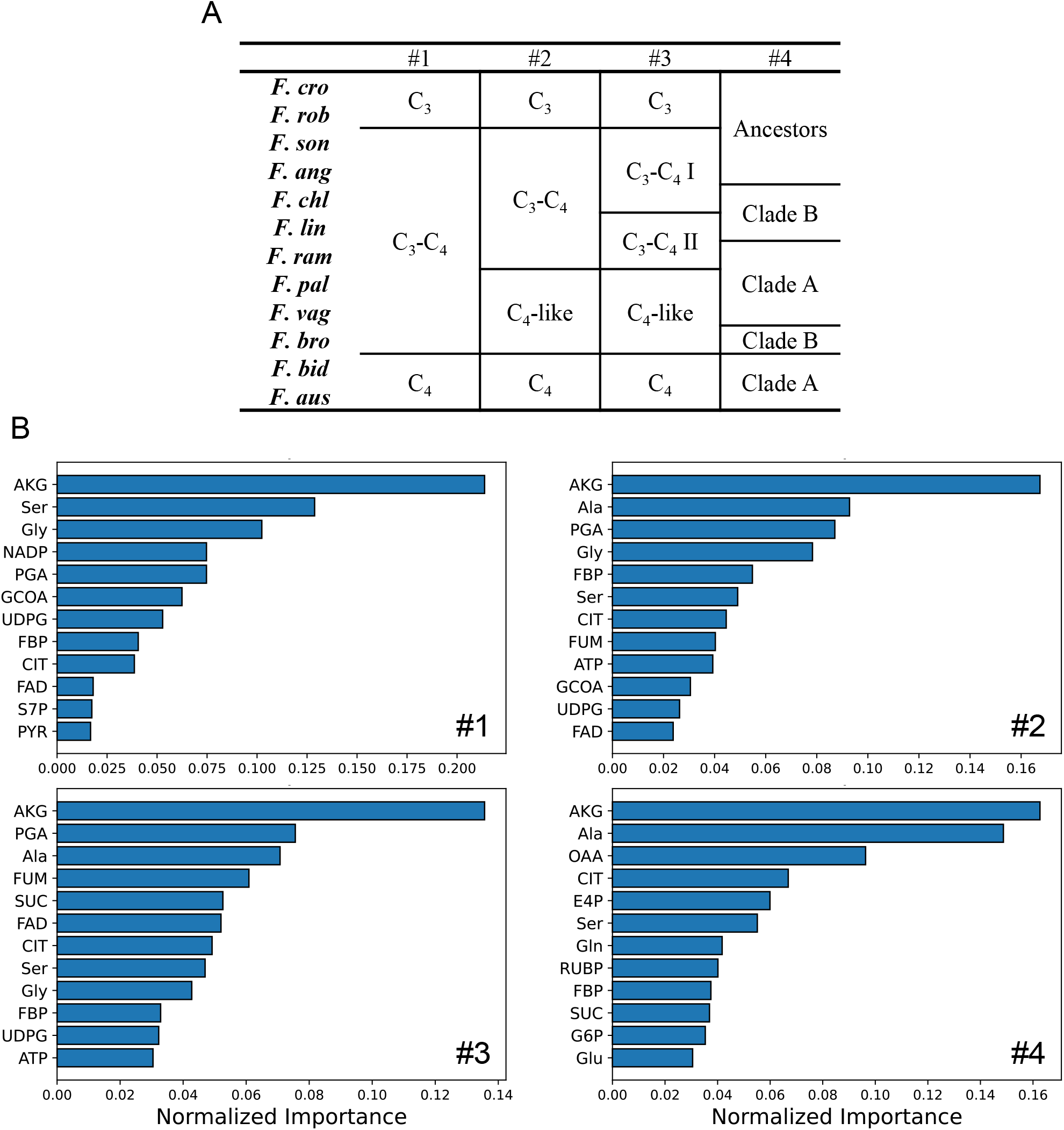
Feature selection analysis of metabolic data. A) Four different schemes for prior grouping. B) important metabolites in each scheme with a cutoff of 80% cumulative importance

**Figure S2.**
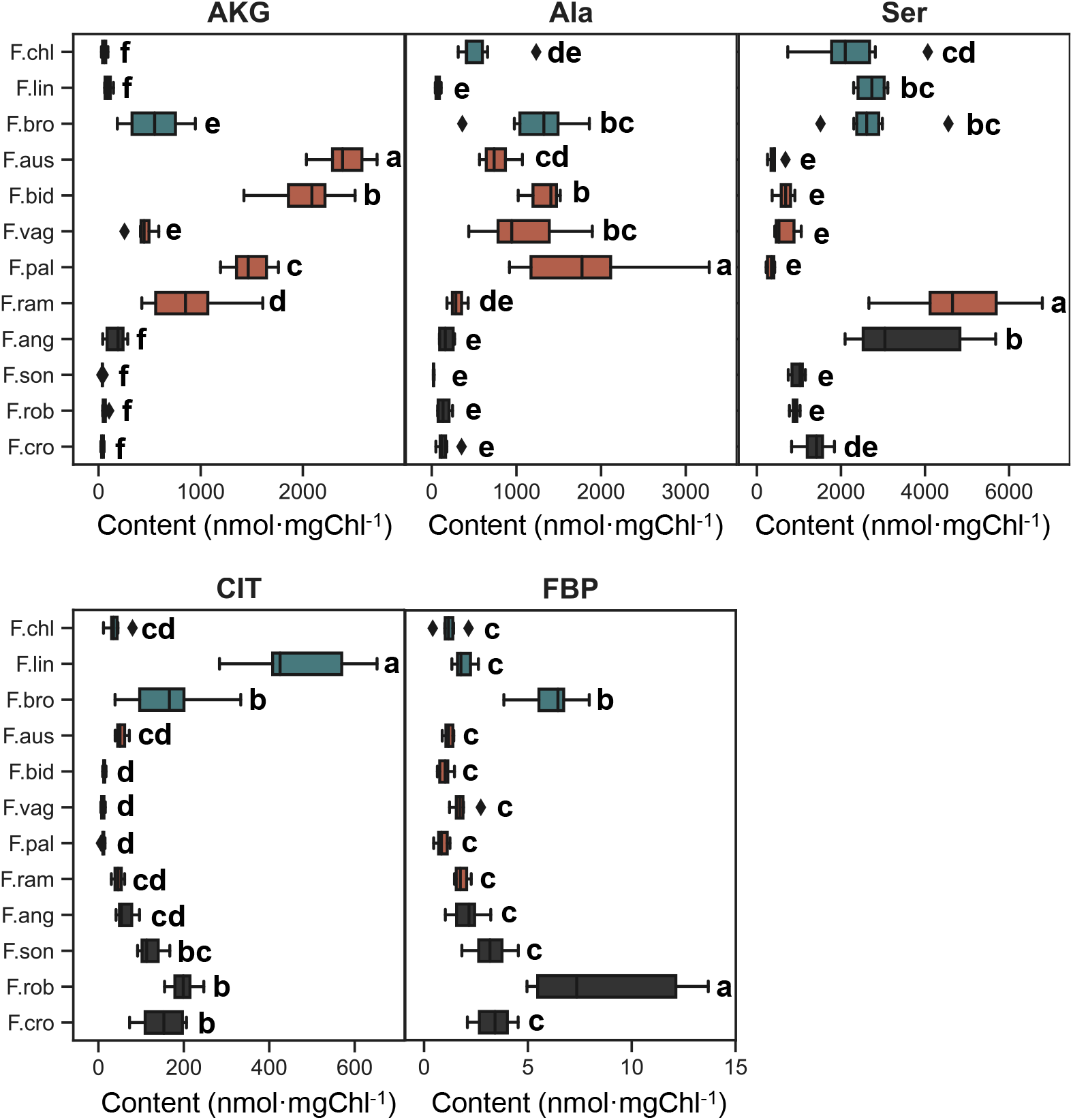
Box plot of five important metabolites. Box with black background indicate the ancestor species in phylogenetic tree, orange indicate the species belong clade A in phylogenetic tree, and green indicate the species belong clade B in phylogenetic tree. Different letters indicate a significant difference between the medians (Kruskal-Wallis tests, α = 0.05, p-values adjusted using false discovery rate) (Benjamini and Hochberg, 1995). n = 6.

**Figure S3.**
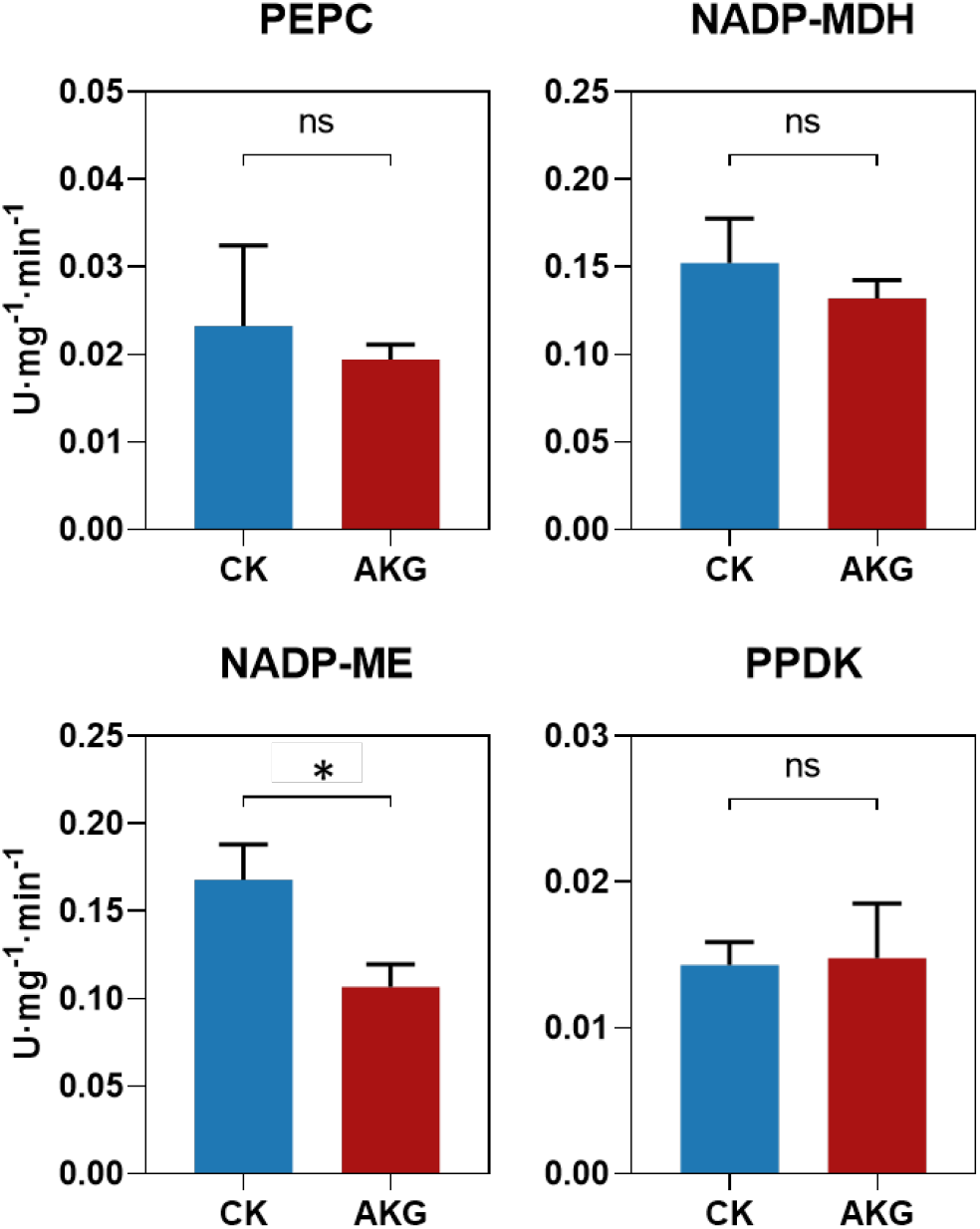
Enzyme activities of C4 photosynthesis in AKG-treated F. ram. Error bar indicates S.D., n = 4. ns.: not significant, *: p < 0.05 (Student’s t-test).

